# A Lipocalin and a Hedgehog-related protein are partners in the *C. elegans* pre-cuticle apical extracellular matrix

**DOI:** 10.64898/2026.07.18.739337

**Authors:** Nicholas D. Serra, Jason Chen, Susanna K. Birnbaum, Sage G. Aviles, Meera V. Sundaram

## Abstract

Apical extracellular matrices (aECMs) line exposed body surfaces to shape tissues and protect them from the environment. These aECMs often organize into complex patterns and structures, but how such matrices assemble remains poorly understood. *Caenorhabditis elegans* cuticle patterns initiate within the transient pre-cuticle, which then helps direct the placement of cuticle collagens. Pre-cuticle patterns arise through post-secretory sorting, which must involve specific molecular interactions among them. Consistent with such a model, Alphafold3 predicts a high confidence physical interaction between two pre-cuticle proteins, the lipocalin LPR-3 and the Hedgehog-related protein WRT-10, with a conserved N-terminal region of LPR-3 forming a β-strand that incorporates into the β-barrel-like structure of the WRT-10 WRT domain. Genetic studies showed that WRT-10 requires this LPR-3 region in order to become properly patterned in the pre-cuticle matrix. Furthermore, WRT-10 and the LPR-3 β-strand region are required to pattern a specific cuticle substructure, the lateral alae ridges, but not for other LPR-3-dependent matrix roles. These data indicate that LPR-3 and WRT-10 are functional partners and support a “landing pad” model whereby physical interactions between them allow LPR-3 to recruit WRT-10 to specific aECM regions. Similar mechanisms may explain how other members of the *C. elegans* Hh-r family associate with the aECM.

**Graphical Abstract:** 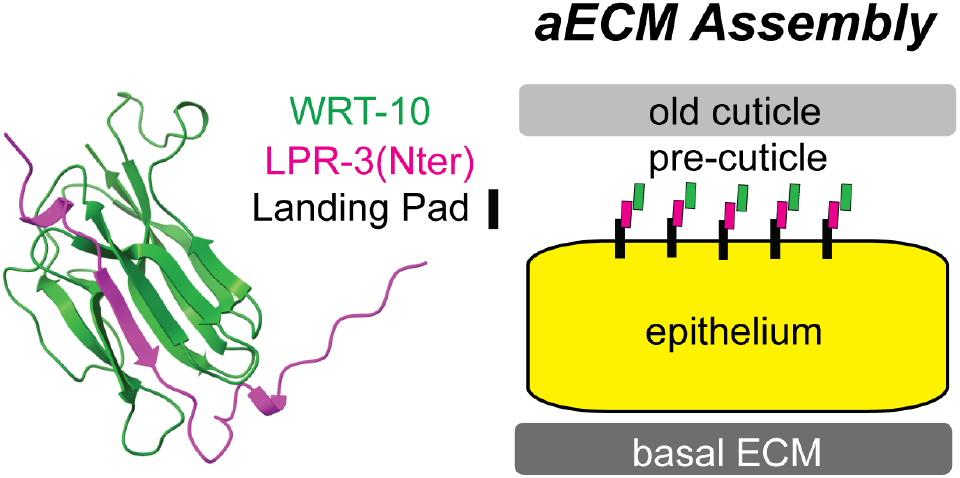

**Article Summary:** All animal skin is covered by a set of proteins, sugars and lipids that comprise the apical extracellular matrix (aECM). These matrix components can be organized into patterned ridges and other distinctive structures. This study addresses how such patterns form in the developing cuticle of the nematode *C. elegans*. The study provides evidence for a regulatory mechanism that enables one matrix protein to establish a pattern and then recruit a second protein into the same pattern.

## Introduction

Apical extracellular matrices (aECMs) coat and protect exposed body surfaces and determine how organisms cope with challenges in their environment (Heiman and Sundaram 2026). These matrices are composed of multiple proteins, glycans and lipids, with their precise composition varying among different tissues and organisms. In many cases, aECMs form multiple layers and other three-dimensional substructures such as pores, ridges, pillars, denticles, or scales (Sapio et al. 2005; Fernandes et al. 2010; Adams et al. 2023; Totz et al. 2024; Itakura et al. 2026). How such precisely patterned matrices assemble in the extracellular environment remains poorly understood.

The nematode *C. elegans* has a collagen-rich cuticle that provides a useful model for studying aECM assembly and function (Sundaram and Pujol 2024). The body cuticle is organized into discrete layers and substructures such as circumferential annuli and furrows and longitudinal ridges (alae) (**Fig. 1A**), each of which contains different collagens (Ragle et al. 2025). Interfacial tubes such as the vulva (**Fig. 1A**) are also lined with a distinct set of cuticle collagens (Ragle et al. 2025). An important insight into cuticle assembly was the discovery of a set of transient “pre-cuticle” matrix proteins that help to pattern the cuticle collagens into specific substructures (Gill et al. 2016; Cohen, Sparacio, et al. 2020; Katz et al. 2022; Sundaram and Pujol 2024; Mazzoli et al. 2026). Although pre-cuticle proteins are not part of the mature cuticle structure, their loss can greatly disrupt cuticle organization.

**Fig. 1.**
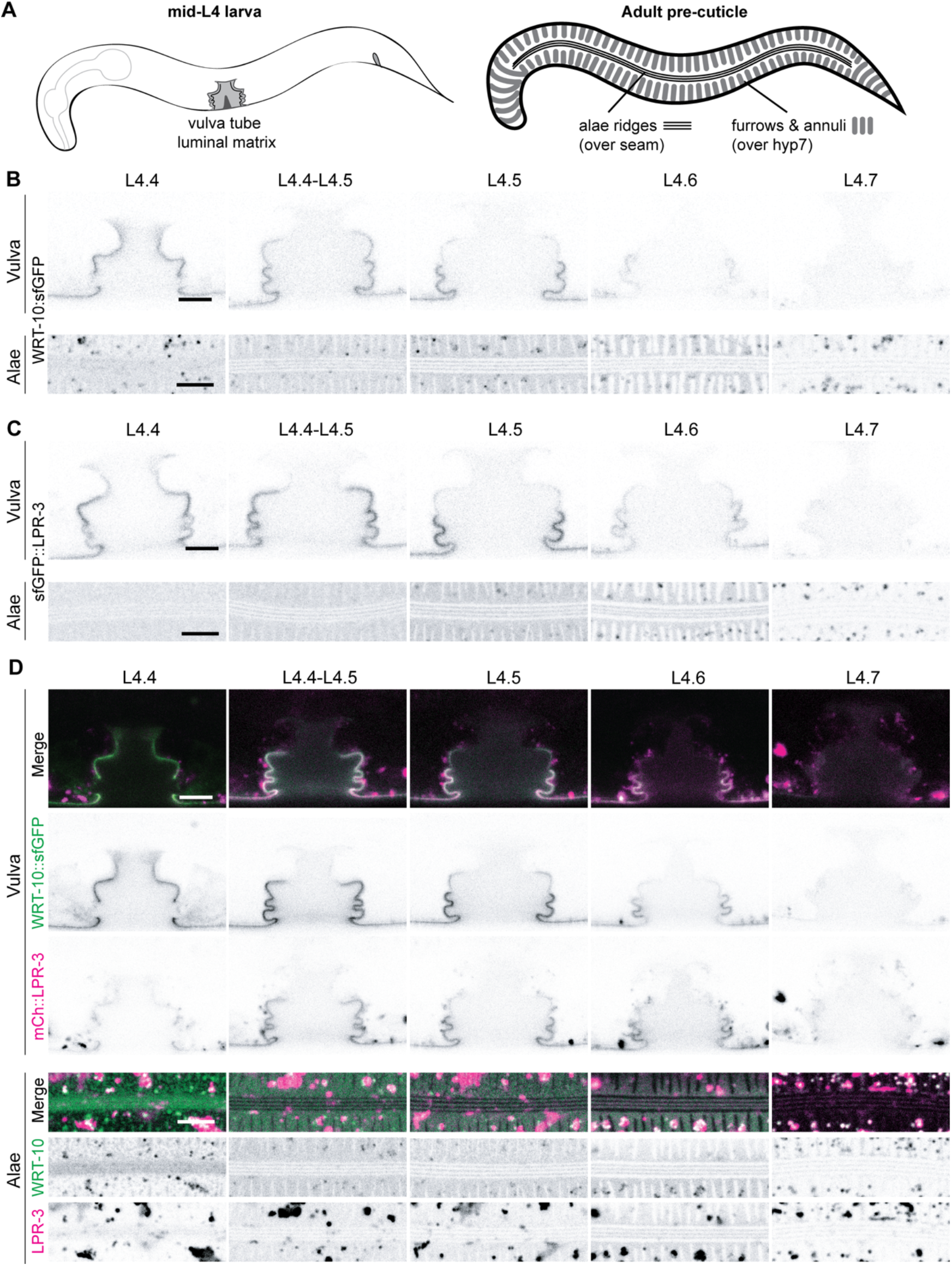
WRT-10 and LPR-3 have similar spatiotemporal dynamics in the aECM. **A.** L4 worm cartoons showing developing vulva tube with luminal matrix (left) and the epidermal pre-cuticle (right) whose patterns predict final adult cuticle substructures. Longitudinal ridges or alae form over the lateral seam epidermis. while circumferential annuli and furrows form over the major epidermis (hyp7). **B-D.** Single confocal Z-slices of the developing vulva luminal matrix and 3 slice projections of developing alae matrix across different mid-L4 stages. Individual channels are shown in inverted grayscale for clarity. All images are representative of at least n=6 per stage. Scale bars, 5 microns. **B.** WRT-10::sfGFP shows dynamic patterns, marking different subsets of vulva cells at each stage and peaking within the alae ridges at the L4.5 and L4.6 stages. **C.** sfGFP::LPR-3 shows similar dynamic patterns to B. **D.** WRT-10::sfGFP and mCh::LPR-3 fusions show substantial overlap across all L4 stages. Note that mCherry is slower to mature than sfGFP (Balleza et al. 2018), which is apparent when comparing LPR-3 fusions across panels C and D at the L4.4 stage.

Pre-cuticle proteins include members of conserved matrix protein families such as zona pellucida (ZP) domain proteins (Kelley et al. 2015; Gill et al. 2016; Vuong-Brender et al. 2017 Jan 1) and extracellular leucine-rich repeat only (eLRRon) proteins (Mancuso et al. 2012), as well as putative lipid transporters (lipocalins) (Forman-Rubinsky et al. 2017; Belfi et al. 2026) and various nematode-specific proteins (Sundaram and Pujol 2024; Sonntag et al. 2025; Mazzoli et al. 2026). Among the latter are several divergent families of Hedgehog-related (Hh-r) proteins, which appear to have both matrix and signaling roles (Burglin and Kuwabara 2006; Hao et al. 2006; Kume et al. 2019; Templeman et al. 2020; Chiyoda et al. 2021; Fung et al. 2023; Shi and Murphy 2023; Serra et al. 2024). Many pre-cuticle proteins are broadly expressed and secreted, but once in the extracellular environment, they assemble into matrix structures only on specific cell surfaces or in specific periodic patterns above the epidermis, providing a pre-pattern that helps guide subsequent collagen addition (Cohen, Sparacio, et al. 2020; Mazzoli et al. 2026). For example, many pre-cuticle factors assemble into longitudinal bands above the lateral “seam” epidermis and promote proper formation of cuticle alae ridges in this location (Katz et al. 2022). The mechanisms that initially pattern pre-cuticle proteins are still largely unknown but are proposed to involve both mechanical cues from the cytoskeleton and molecular sorting that relies on specific interactions among them (Gill et al. 2016; Low et al. 2019; Cohen, Bermudez, et al. 2020; Katz et al. 2022; Schmidt et al. 2025; Heiman and Sundaram 2026; Mazzoli et al. 2026; Sarwar et al. 2026).

Here we provide evidence that a lipocalin, LPR-3, and a Hh-r protein, WRT-10, are partners in the pre-cuticle aECM and physically interact to pattern the lateral alae. LPR-3 plays many important matrix-related roles, being essential for embryonic hatching, protection of narrow tube integrity, molting, establishment of cuticle barrier function, and patterning of the cuticle alae ridges (Forman-Rubinsky et al. 2017; Katz et al. 2022). WRT-10 has more limited functional requirements, being required only for patterning of the cuticle alae ridges (Serra et al. 2024). In addition to their matrix roles, both proteins have also been implicated in signaling processes (Templeman et al. 2020; Shi and Murphy 2023; Wu et al. 2025). Through imaging of endogenous fluorescent protein fusions, we find that WRT-10 and LPR-3 show very similar spatiotemporal patterns of expression and localization in L4 larvae, during the period of adult cuticle construction. Furthermore, Alphafold3 (Abramson et al. 2024) predicts, with high confidence, a physical interaction between a unique region in the LPR-3 N-terminus and the WRT-10 Warthog (WRT) domain; this prediction extends to distant orthologs in other nematodes such as the parasites *Strongyloides ratti* and *Brugia malayi*. Deletion of the *C. elegans* LPR-3 N-terminal domain mimics deletion of WRT-10, causing specific alae ridge defects, failure to properly pattern both LPR-3 and WRT-10 within the alae matrix, and diminished co-localization of the two proteins across other aECM regions. These genetic data support the Alphafold3 predictions and show that LPR-3 and WRT-10 are functional partners in the pre-cuticle aECM, with LPR-3 serving as a molecular “landing pad” that recruits WRT-10 to specific matrix regions.

## Materials and Methods

### Strains and Animal husbandry

*C. elegans* were maintained at 20°C on NGM plates seeded with OP50-1, as described (Brenner 1974). Strains used are listed in S1 Table. Information from Wormbase and the Alliance of Genome Resources (Sternberg et al. 2024) were used for all aspects of experimental design.

To generate *lpr-3(cs266 cs315 [mCherry::LPR-3(6.Nter)])*, CRISPR/Cas9-mediated genome editing was performed on strain UP3808 using methods as in (Dokshin et al. 2018). Purified Cas9 was obtained from University of California Berkeley. The guide RNAs used were a mix of tracrRNA, Ce.Cas9.LPR-3.1.AK (GCTGGGAGATGAATTACTAC) and Ce.Cas9.LPR-3.1.AS (TTGGACGGTGGTGTTGTGCT), and the repair template was single stranded oligonucleotide 5’AATTGTATAAGGGTGCTATTAGCGAAGCAGACGTAGCACAACACCACCGTCCAAGATTTTTAGGACAGGT3’ (all from IDT). F1 and F2 candidates were screened using PCR primers oNS40 CAC CTC CCA CAA CGA GGA TTAC (in mCherry) and oNS38 ATC TTG GAC GGT GGT GTT GTG (in *lpr-3*) and lesions confirmed by sequencing. The strain was outcrossed to N2 four times (generating strain UP4435) prior to characterization. Rescue experiments used a previously described fosmid-based *lpr-3+* transgene (*csEx436*) (Forman-Rubinsky et al. 2017).

### Alphafold analyses

Alphafold3 (Abramson et al. 2024) was used for all pairwise analyses. Predicted structures from the top scoring models (”model 0”) are shown. Visual representations were created using UCSF ChimeraX (Pettersen et al. 2021).

### Microscopy

Differential Interference Contrast (DIC) and BLI-6::mNG epifluorescent images were collected with a Zeiss Axioskop (Carl Zeiss Microscopy) and a Leica DFC360 FX camera. Confocal images were collected with a Leica Dmi8 microscope using an HC PL APO 63x objective (Numerical Aperture 1.3), 600 Hz scanning speed, 1X line accumulation, 1X line average, and a pinhole setting of 1 AU. SfGFP fusions were excited with a 488 nm laser and emissions between 493 and 547 nm detected using a HyD sensor. mCherry fusion proteins were excited with a 552 nm laser and emissions between 583 and 784 nm detected by a HyD sensor. Laser power was set at 3% and gain settings varied (50-90%) based on fusion protein. Unless otherwise indicated, all confocal images shown are single Z-slices.

Larvae were immobilized with 10 mM levamisole in M9 buffer and mounted on a 5% agarose pad; for confocal imaging, pads were supplemented with 20mM sodium azide. L4 stages were assessed based on vulva lumen morphology (Mok et al. 2015; Cohen, Sparacio, et al. 2020).

### Image quantification and statistical analyses

Alae defects were quantified using 120 micron-wide images of BLI-6::mNG patterns in 24-hour adults. Three images (from the anterior, mid-body, and posterior regions) were taken per animal and then scored by a researcher blinded to genotypes. The four categories are: 1. *Normal*: Consistent fluorescent signal with no interruptions; 2. *Minor:* Less than three alae breaks or ten incomplete breaks, defined as nicks in the ridges and dips in band intensity; 3. *Partially Disorganized*: Four or more alae breaks interspersed with regions of normality, with at least half of the image appearing normal; 4. *Disorganized*: Alae breaks throughout the length of the image. For statistical analysis, categories 1 and 2 were combined and categories 3 and 4 were combined so that outcomes were classified as ∼normal vs abnormal for the Fisher’s Exact Test, which was carried out using GraphPad Prism. Identical results were obtained when category 2 instead was combined with the abnormal categories 3 and 4.

Peak fluorescence intensity across the epidermis was measured using a 10-point wide line and the Plot Profile Tool in FIJI (Schindelin et al. 2012); the line was drawn circumferentially across the entire width of the seam at the image midpoint (to capture the 3 alae ridges) or longitudinally across a 10 micron length of hyp7 (to capture 6-8 annuli but avoid intracellular puncta). For each tissue, peak values were averaged to obtain one seam and one hyp data point per animal. For Fig. 5E, Mander’s overlap coefficients were calculated using the JaCoP plug-in in FIJI (Bolte and Cordelières 2006). For statistical analyses, data were compared using a non-parametric Mann-Whitney test.

### Data availability

All data are available in the figures and supplements. S1 Fig. and S2 Table show additional Alphafold3 data and S3-S9 Tables contain the raw data used for graphs. Strains (S1 Table) are available from the Caenorhabditis Genetics Center (U. Minnesota) or from the corresponding author.

## Results

### WRT-10 and LPR-3 fusions show similar patterns of spatiotemporal expression

Nematodes have ∼60 Hh-r proteins that are thought to be evolutionarily related to the canonical Hh signaling ligand, but which have diverged significantly in sequence and contain nematode-specific cysteine repeat domains (WRT, GRL, GRD, QUA) (Burglin and Kuwabara 2006; Bürglin 2008). Protein structure databases predict that Hh-r proteins are secreted and that each of the Hh-r domains forms a characteristic compact fold (Senior et al. 2020; Abramson et al. 2024). Recent studies demonstrated that several Hh-r proteins are stably associated with specific regions of the cuticle or pre-cuticle (Chiyoda et al. 2021; Fung et al. 2023; Serra et al. 2024). Among these is WRT-10, which we recently showed is a pre-cuticle factor important for adult alae ridge patterning (Serra et al. 2024). This raised the question of how WRT-10 associates with and modifies the aECM.

When imaging an endogenous WRT-10::sfGFP fusion, we noticed that its spatiotemporal pattern of matrix localization was strikingly similar to what we had previously reported for the lipocalin LPR-3 (Forman-Rubinsky et al. 2017; Cohen, Sparacio, et al. 2020; Katz et al. 2022) (**Fig. 1**). For example, in the developing vulva tube lumen, WRT-10::sfGFP and sfGFP::LPR-3 fusions both showed the same dynamic pattern of localization to different vulva cell surfaces: Each initially associated with all vulva cell apical surfaces and then was progressively lost from more dorsal cell surfaces before being entirely cleared (**Fig. 1A-C**). Similarly, in the epidermal pre-cuticle, both fusions were secreted broadly, then patterned to label developing annuli and alae ridges, and then cleared by endocytosis at the same timepoints (**Fig. 1A-C**). Imaging of a double labelled strain containing both WRT-10::sfGFP and mCherry::LPR-3 confirmed substantial overlap across these tissues and timepoints (**Fig. 1D**).

Importantly, while shared between WRT-10 and LPR-3, these specific patterns and their timing are different than those reported for other pre-cuticle fusions (Gill et al. 2016; Cohen, Sparacio, et al. 2020; Katz et al. 2022). These data suggest that WRT-10 and LPR-3 might function together within the pre-cuticle.

### Alphafold3 predicts that WRT-10 and LPR-3 can physically interact via a β-strand addition mechanism

Consistent with the idea that WRT-10 and LPR-3 act together, the protein structure prediction program Alphafold3 (Abramson et al. 2024) predicted that WRT-10 and LPR-3 could physically interact (**Fig. 2**). In the Alphafold3 models, the WRT-10 WRT domain appears as a β-sandwich-like structure consisting of eight β-strands, while the WRT-10 C-terminal region appears largely unstructured (**Fig. 2A,B; S1 Fig.**). Conversely, the N-terminal region of LPR-3 appears largely unstructured, while the lipocalin domain forms the signature lipocalin cup, also consisting of eight β-strands (Flower et al. 2000) (**Fig. 2A,B; S1 Fig.**). In tests of the full-length proteins (minus their signal peptides), Alphafold3 predicted an interaction between the WRT-10 WRT domain and a small region of the extreme LPR-3 N-terminus, which formed an extra β-strand specifically in the presence of WRT-10 (**Fig. 2B**). Although the interface predicted template modeling (IpTM) score of 0.62 was in the “gray zone” for confidence, the relevant region of the structure was in the higher confidence range (>90) of pLDDT (Local Difference Distance Test) scores (**Fig. 2B**). Because IpTM scores would necessarily be reduced by the presence of the long unstructured regions in both proteins, we further tested the isolated WRT-10 WRT domain (residues 28-154) with LPR-3 N-terminal residues 18-50; this focused test yielded a high confidence IpTM score of 0.85 (**Fig. 2B**), suggesting that WRT-10 and LPR-3 are likely to interact *in vivo*.

**Fig. 2.**
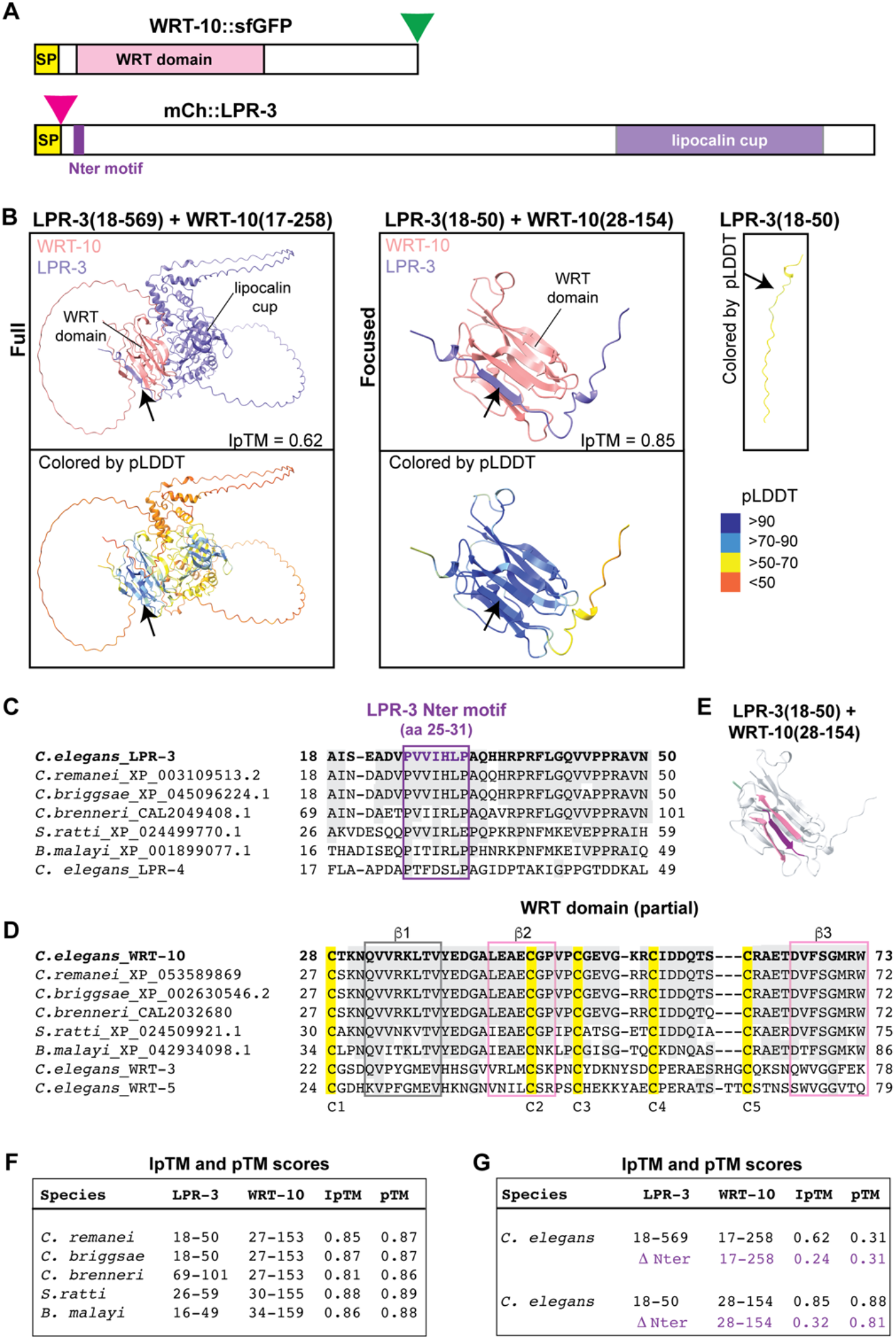
Alphafold predicts an interaction between WRT-10 and the LPR-3 N-terminus. **A.** Schematics of the WRT-10 and LPR-3 proteins, with domains and tag locations indicated. SP, signal peptide. Green triangle, sfGFP. Magenta triangle, mCherry. **B.** Alphafold prediction models showing interactions of the full-length proteins after signal peptide removal (left), a focused view of the interacting regions (middle), and the isolated LPR-3 N-terminal region (right). **C.** LPR-3 partial sequence alignment with other nematode orthologs and the closest paralog LPR-4. Purple box indicates the Nter motif predicted to interact with WRT-10. Grey highlights residues identical between LPR-3 and the other proteins. **D.** WRT-10 partial sequence alignment with other nematode orthologs and closest paralogs WRT-3 and WRT-5. Yellow highlights conserved cysteines and boxes indicate predicted β-strands in the WRT domain. Grey highlights residues identical between WRT-10 and the other proteins. **E.** Schematic of LPR-3-WRT-10 interacting regions with predicted β-strands colored as in C,D. **F.** Table of iPTM values showing evolutionary conservation of the predicted interaction. See panels C and D for the accession numbers of sequences used for this analysis. **G.** Table of iPTM values showing effect of deleting the 7 amino acid LPR-3(Nter) motif.

The predicted interacting regions of LPR-3 and WRT-10 were highly conserved in orthologs from other *Caenorhabditis* species (>75% identity) and moderately conserved in other more distantly-related nematodes such as the parasites *Strongyloides ratti* and *Brugia malayi* (>40% identity) (**Fig. 2C-E**). Strikingly, even in the more distantly related nematodes, these regions of LPR-3 and WRT-10 were predicted to interact with high confidence (**Fig. 2F**). In contrast, other *C. elegans* lipocalins and WRT proteins have more limited similarity to LPR-3 or WRT-10 across these regions (**Fig. 2C,D**) and were not predicted to interact with them (**S2 Table).** Together, these analyses suggest that the predicted interaction between LPR-3 and WRT-10 is likely to be functionally important and specific.

### Deletion of the LPR-3 Nter motif phenocopies *wrt-10(-)* alae patterning defects

To test the importance of a possible direct interaction involving the LPR-3 N-terminal domain, we used CRISPR/Cas9 to delete seven amino acids (PVVIHLP) from this region by modifying the endogenous *lpr-3* locus. These residues encompass the predicted β-strand that interacts with the WRT domain in the Alphafold3 models (**Fig. 2B,C,E**), and the LPR-3(6.Nter) deletion was predicted to severely reduce or eliminate the interaction (**Fig. 2G**). The deletion was made within our existing mCh::LPR-3 fusion strain, so that the localization of the mutant protein could be assessed. *lpr-3(cs315)* mutants (hereafter called *lpr-3(6.Nter)*) were fully viable and healthy (247/248 live), without any sign of the lethal phenotypes previously described for *lpr-3* mutants nor of the molting defective phenotypes of *lpr-3(RNAi)* treated animals (Forman-Rubinsky et al. 2017). However, the mutants did show alae defects similar to those of *wrt-10(aus36)* deletion mutants (hereafter *wrt-10(-)*), as observed by differential interference contrast (DIC) microscopy (**Fig. 3A**) or with a fluorescently-tagged alae collagen, BLI-6 (Adams et al. 2023) (**Fig. 3B**). These *lpr-3(6.Nter)* defects were rescued by an *lpr-3+* transgene (**Fig. 3A-C**). Notably, in both *lpr-3(6.Nter)* and *wrt-10(-)* mutants, alae defects predominantly (but not exclusively) affected the middle alae ridge and were more severe in the posterior body region than in the anterior or mid-body (**Fig. 3C**). The reasons for these specific aspects of the phenotype are not obvious, given that in wild type both proteins localize equally to (and along the full length of) all three developing ridges (**Fig. 1**). Importantly, *wrt-10(-); lpr-3(6.Nter)* double mutants showed alae defects that were no more severe than those of single mutants, consistent with WRT-10 and LPR-3 acting together (**Fig. 3A-C**).

**Fig. 3.**
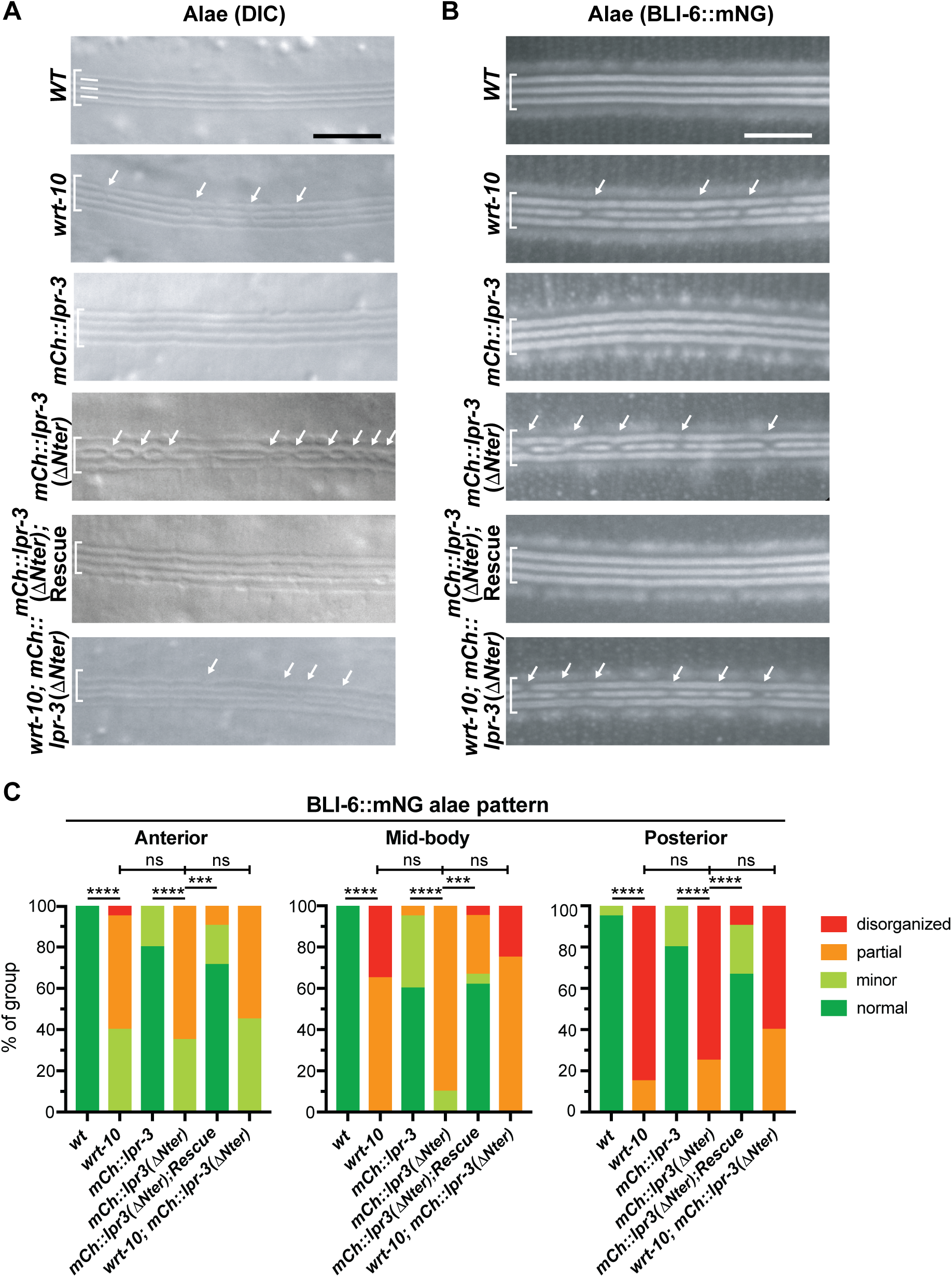
*lpr-3(6.Nter)* phenocopies *wrt-10(-)* mutant alae defects. **A.** DIC images of alae in young (non-gravid) adults. **B.** Epifluorescent images of alae collagen BLI-6::mNG (Adams et al. 2023) in 24 hr adults. Brackets indicate alae ridge positions and arrows indicate alae ridge interruptions. Scale bars, 5 microns. **C.** Severity of alae defects varies across different body regions. Defects were quantified based on the BLI-6::mNG pattern, since the mutant alae often appeared faint and difficult to assess by DIC. ***p<0.001, ****p<0.0001, Fisher’s Exact Test (normal vs. abnormal). See Materials and Methods for details.

### Deletion of the LPR-3 Nter motif or loss of WRT-10 cause similar seam-specific changes in the localization of LPR-3 fusions

Examination of the mutant mCh::LPR-3(6.Nter) fusion revealed that it still marked the three developing alae ridges, but in an irregular fashion with various small gaps, consistent with perturbed development of all three ridges (**Fig. 4A**). mCh::LPR-3(6.Nter) was also dimmer across the seam at the L4.6 stage **(Fig. 4A,B)**, suggesting premature loss from the matrix. However, the fusion remained normal in the epidermal and vulva pre-cuticles (**S2 Fig.**). Very similar seam-specific changes in sfGFP::LPR-3 were observed in *wrt-10(-)* mutants (**Fig. 4C,D and S2 Fig.**). These results indicate that WRT-10 specifically assists in LPR-3 alae matrix patterning and retention and strengthen the conclusion that deleting the LPR-3 N-terminal domain causes phenotypes similar to loss of WRT-10, without disrupting other LPR-3 functions.

**Fig. 4.**
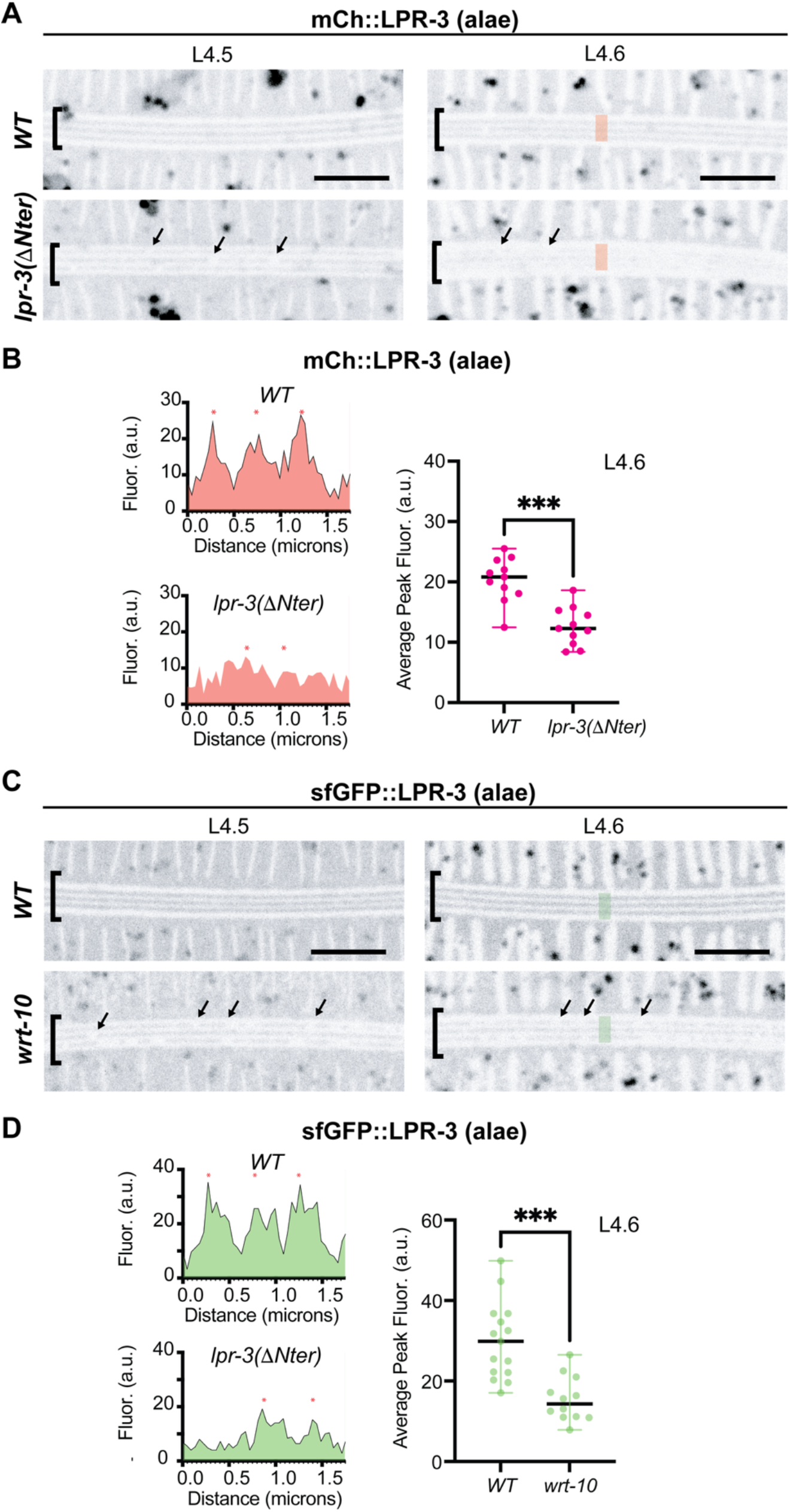
*lpr-3(6.Nter)* and *wrt-10(-)* mutants disrupt LPR-3 patterning in developing alae. **A,C.** mCh::LPR-3 (A) or sfGFP::LPR-3 (C) in developing alae at the indicated stages. Images are maximum intensity projections of three confocal z-slices, shown in inverted grayscale. The alae signal (brackets) is interrupted (arrows) in *lpr-3(6.Nter)* and *wrt-10(aus36)* mutants and signal intensities are diminished. Colored bars indicate method for quantifying fluorescence intensity in B,D. Scale bars, 5 microns. **B,D.** Quantifications of peak fluorescent signal intensity across the L4.6 seam, as measured using the FIJI Plot Profile tool (n>10 for each genotype). Example line scans are shown at left and asterisks indicate peak regions called. a.u., arbitrary units. See Materials and Methods for additional details. ***, p<0.001, Mann-Whitney U test.

### Deletion of the LPR-3 Nter motif broadly disrupts WRT-10 patterning and LPR-3-WRT-10 colocalization

Finally, we observed that LPR-3(6.Nter) dramatically affected WRT-10::sfGFP matrix localization throughout the animal (**Fig. 5**), indicating that LPR-3 plays a key role in WRT-10 matrix patterning. Over the seam epidermis, WRT-10::sfGFP signal failed to resolve into discrete alae ridges, even in animals where the mCh::LPR-3 pattern remained largely intact; instead, WRT-10::sfGFP appeared in a general dim haze (**Fig. 5A,C**). The WRT-10::sfGFP signal was also much reduced in the epidermal annuli and less efficiently excluded from furrow regions, although mCh::LPR-3 had normal patterning (**Fig. 5B,C**). Equally striking was the localization pattern in the vulva, where WRT-10::sfGFP now partitioned away from mCh::LPR-3 to label different vulva cell surfaces (**Fig. 5D,E**), suggesting the existence of an alternative binding partner in that region. These data indicate that the normally observed co-localization of WRT-10 and LPR-3 depends on the LPR-3 N-terminal motif and are consistent with the two proteins interacting via this motif.

**Fig. 5.**
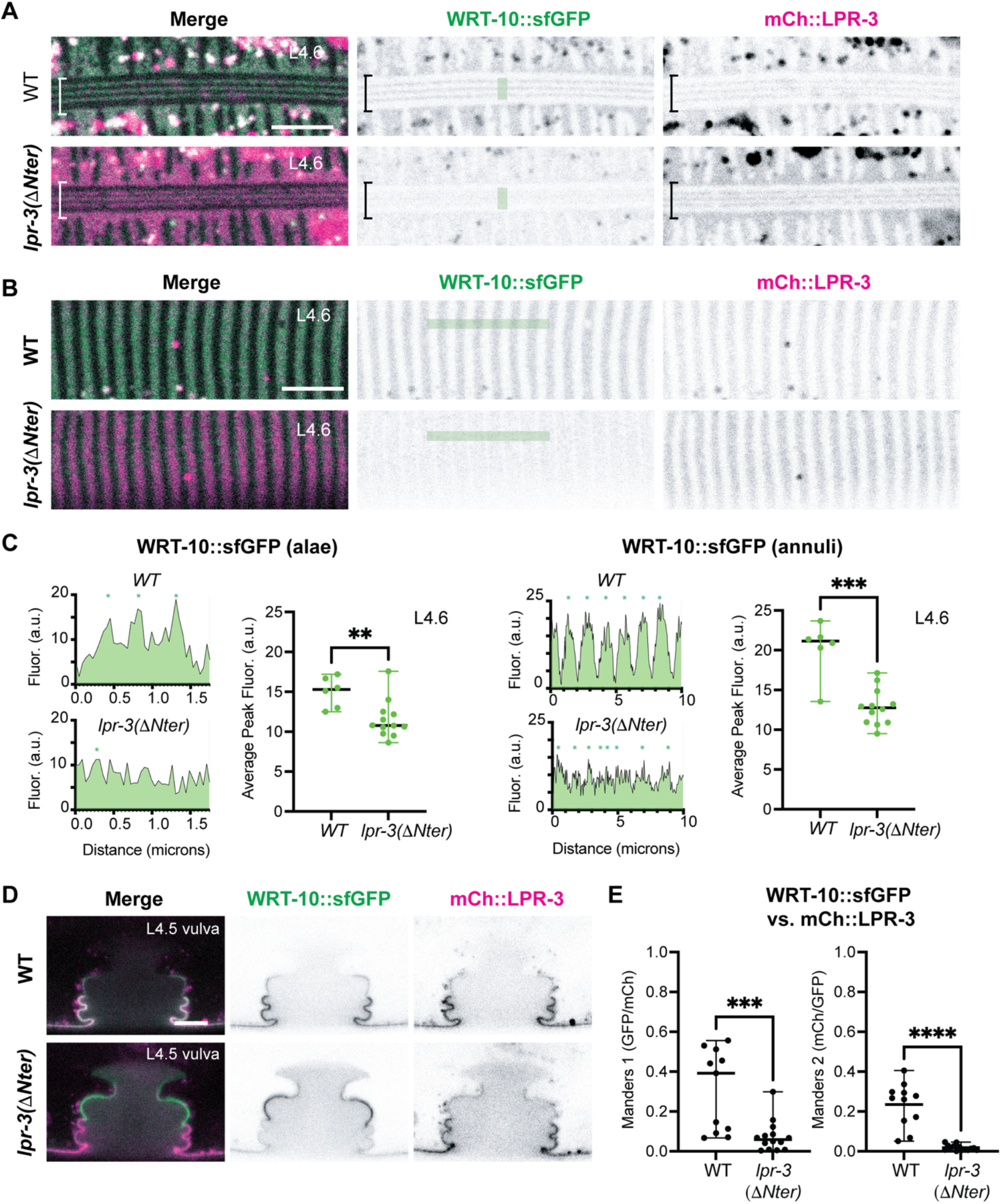
*lpr-3(6.Nter)* broadly disrupts WRT-10 matrix patterning and WRT-10-LPR-3 colocalization. **A.** Alae localization of WRT-10::SfGFP and mCh::LPR-3. In *lpr-3(6.Nter)* mutants, the WRT-10 signal is incompletely patterned at L4.5 and then decays by L4.6. **B.** Annuli localization of WRT-10::SfGFP and mCh::LPR-3. In *lpr-3(6.Nter)* mutants, the WRT-10 signal is weak and no longer coincides precisely with the LPR-3 signal. Colored bars indicate method for quantifying fluorescence intensity in C. **C.** Quantification of peak WRT-10::sfGFP fluorescence intensity in the alae (left) and annuli (right) as measured using the FIJI Plot Profile tool (see Materials and Methods). Example plots of line scans illustrate the lack of patterning. a.u., arbitrary units. **D.** Vulva localization of WRT-10::SfGFP and mCh::LPR-3. In *lpr-3(6.Nter)* mutants, the two proteins no longer localize to the same cell surfaces. **E.** Mander’s co-efficients of overlap are greatly reduced in *lpr-3(6.Nter)* mutants. ** p<0.01, *** p<0.001, **** p<0.0001, Mann-Whitney U test. All images representative of at least n=10 images. Note that only n=6 WT L4.6 stage animals could be included in the quantifications because the other n=6 images were taken with different settings; however, all appeared similarly patterned, unlike the mutants. Scale bars 5 microns.

## Discussion

Many of the elaborate patterns observed in the *C. elegans* cuticle initiate within the transient pre-cuticle, which then helps guide appropriate placement of collagens in the mature cuticle structure. Pre-cuticle patterning involves post-secretory mechanisms that must involve specific molecular interactions among its components. Here we provided genetic evidence that two specific pre-cuticle proteins, the lipocalin LPR-3 and the Hh-r protein WRT-10, are functional interacting partners, with LPR-3 serving as a key determinant of WRT-10 matrix patterning. These studies provide molecular insight into how an Hh-r protein incorporates into the aECM and suggest a β-strand addition interaction model that could be relevant to other members of this family. Furthermore, the results support the previously proposed “landing pad” model for aECM patterning **(Fig. 6)** (Heiman and Sundaram 2026) but suggest that pre-cuticle factors have multiple binding partners whose relative affinity or availability ultimately determine cell-type specific and substructure-specific matrix assembly patterns.

**Fig. 6.**
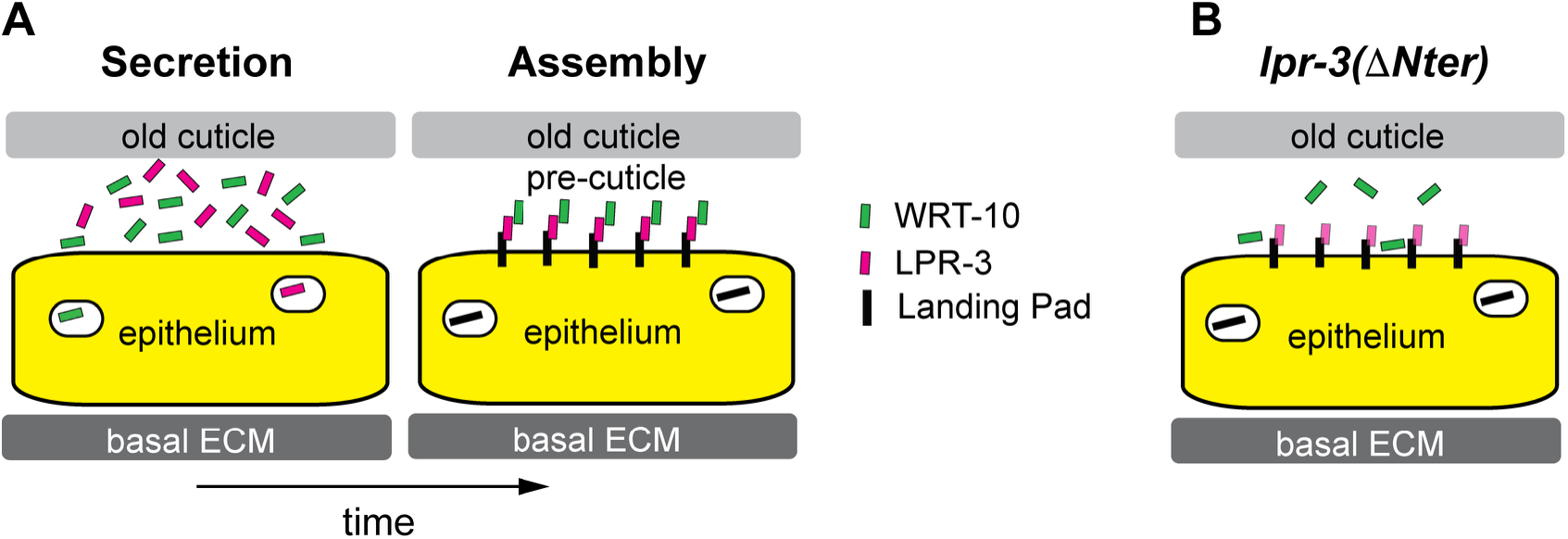
Landing Pad Model for LPR-3 and WRT-10 matrix assembly. **A.** Under normal conditions, pre-cuticle proteins such as LPR-3 and WRT-10 are initially secreted in soluble form into the space below the old cuticle, which helps confine them near the cell surface. Once the cell displays an appropriate landing pad (black rectangle), it recruits LPR-3 (magenta), which in turn recruits WRT-10 (green) for matrix assembly. **B.** In *lpr-3(6.Nter)* mutants, LPR-3 can still assemble at least in part, but WRT-10 is no longer recruited to the same locations.

### LPR-3, WRT-10, and the Landing Pad model of aECM Assembly

Classically, invertebrate cuticles have been thought to develop through a “3D printing”-like process involving sequential secretion and rapid membrane-proximal assembly of new layers (Wigglesworth 1948; Kim et al. 2019; Sundaram and Pujol 2024). However, recent studies in *C. elegans* have challenged this model by showing that multiple pre-cuticle and cuticle proteins are initially broadly secreted and only later assembled into patterned structures, sometimes on cells that are different from where they originated (Gill et al. 2016; Low et al. 2019; Cohen, Sparacio, et al. 2020; Katz et al. 2022; Birnbaum et al. 2023; Ragle et al. 2025; Schmidt et al. 2025; Mazzoli et al. 2026). Such observations suggest that extracellular molecular sorting processes such as phase separation or thermodynamic self-assembly are likely to be important for patterning. We have also proposed the concept of molecular “landing pads” that allow specific cell surfaces to recruit partner aECM proteins from the environment for local assembly (Gill et al. 2016; Low et al. 2019; Schmidt et al. 2025; Heiman and Sundaram 2026). All of these models highlight the need to identify relevant protein-protein interactions in order to understand the mechanisms of aECM patterning.

LPR-3 and WRT-10 are two pre-cuticle proteins that nicely illustrate the problem. Both are broadly expressed and show post-secretory patterning that is highly specific yet dynamic over developmental time (**Fig. 1**). Neither protein has a membrane-spanning domain or predicted GPI anchor, so their association with specific cell surfaces and their aECMs must depend on interactions with other factors, such as cell- or matrix-associated landing pads. The data reported here suggest that as yet unidentified landing pad(s) initially recruit LPR-3 to the matrix, and that LPR-3 then functions as a landing pad for WRT-10, recruiting it to the same matrix regions **(Fig. 6)**. WRT-10 in turn stabilizes LPR-3 within the alae matrix. However, WRT-10 must also have additional matrix partners, since in the absence of LPR-3(Nter) it can still associate with some matrix regions. These observations reveal that matrix assembly is not a simple linear cascade in which each aECM protein has a single primary landing pad; instead, like LPR-3 and WRT-10, many aECM proteins are likely to have multiple potential partners. The dynamic changes in pre-cuticle patterning observed during development likely reflect a changing landscape of partner availability and affinity.

### β-strand addition or removal as a mechanism of aECM protein regulation

Although very different from each other in sequence and specific structure, several families of pre-cuticle proteins, including lipocalins (Flower et al. 2000), Hh-r proteins (Senior et al. 2020), and ZP module proteins (Bokhove and Jovine 2018), all have barrel- or sandwich-like structures composed primarily of β-strands. In ZP module proteins, removal of a C-terminal sequence that comprises the final β-strand in the ZPc immunoglobulin-like fold is a key regulatory step in ZP polymerization for matrix assembly (Jovine et al. 2004). In the interaction model described here, an initially unstructured region of the LPR-3 N-terminus is able to form an extra β-strand that incorporates into the WRT-10 β-sandwich structure, without removal of any of the pre-existing WRT-10 β-strands. Similar β-strand addition strategies underlie many other protein-protein interactions (Remaut and Waksman 2006; Cheng et al. 2013) and could be common among aECM proteins. For example, in *C. elegans,* the Groundhog-like (GRL) domain of the Hh-r protein GRL-18 drives its matrix association with developing cuticle pore structures (Fung et al. 2023) and the ZP-c domain of LET-653 drives its matrix association with specific vulva tube cell surfaces (Cohen, Bermudez, et al. 2020). These domains have been proposed to interact with unknown molecular partners or landing pads on destination surfaces. It will be interesting to identify relevant partners of these other aECM proteins and to determine if any of their interactions are also driven by β-strand addition as suggested here for WRT-10 and LPR-3.

### Matrix vs. Signaling Roles of Lipocalins and Hh-r proteins

In addition to their matrix roles, both LPR-3 and WRT-10 have also been implicated in signaling processes. Lipocalins can bind various lipids, including signaling lipids, within their cup-like structure (Flower et al. 2000). Indeed, like the mammalian lipocalin ApoM, LPR-3 can bind the signaling lipid sphingosine-1-phosphate (S1P), which interacts with the S1P receptor in neurons to affect learning (Wu et al. 2025). Multiple Hh-r proteins have been proposed to act as signaling ligands, possibly via Patched-related receptors, although *C. elegans* lacks a Smoothened ortholog and other components of the canonical Hh signaling pathway (Burglin and Kuwabara 2006; Kume et al. 2019; Chiyoda et al. 2021; Emans et al. 2023). WRT-10 has been implicated in both soma-to-germline and germline-to-soma signaling to affect reproduction during aging (Templeman et al. 2020; Shi and Murphy 2023), although we did not detect obvious reproductive changes in *wrt-10* single mutants **(S3 Fig.).** If and how LPR-3-WRT-10 interactions and matrix association would affect these signaling roles are open questions, but neither LPR-3 nor WRT-10 fusions are visible in adults, when these signaling processes are proposed to occur. Therefore, it is possible that phenotypes attributed to signaling are instead indirect consequences of abnormal matrix development defects in these mutants. Further studies will be needed to disentangle the structural and signaling roles of lipocalins and Hh-r mutants, and to determine how each impacts the other.

## Acknowledgements

We thank David Fay for his tutorials on use of Alphafold and members of our lab and the UPenn worm group for helpful discussions and advice. We specifically thank Yuhao Wan for technical assistance and Prioty Sarwar for her initial insight that the severity of alae defects can vary across body regions. Some strains were obtained from the *Caenorhabditis* Genetics Center (U. Minnesota), which is funded by the NIH Office of Research Infrastructure Programs (P40 OD010440).

## Funding

This work was supported by National Institutes of Health grant R35 GM136315 to M.V.S.

## Conflicts of Interest

The authors declare that they do not have any conflicts of interest.

## Supplemental Information

**S1 Table.** *C. elegans* strains used in this study.

|  |  |  |
| --- | --- | --- |
| S1 Table. | C. elegans strains used in this study |  |
| Strain | Genotype | Source |
| CZ24114 | bli-6(ju1914 [BLI-6::mNG::3xFLAG]) IV | Adams et al 2023 |
| HRN666 | wrt-10(aus36) II | Sherry et al 2019 |
| PHX7022 | wrt-10(syb7022[WRT-10::sfGFP]) II | Serra et al 2024 |
| UP3666 | lpr-3(cs250 [sfGFP::LPR-3]) X | Cohen et al 2020 |
| UP3808 | lpr-3(cs266[mCherry::LPR-3]) X | Aviles et al 2026 |
| UP4255 | wrt-10(syb7022[WRT-10::sfGFP]) II; lpr-3(cs266 [mCherry::LPR-3]) X | This work |
| UP4299 | wrt-10(aus36) II; lpr-3(cs250[ssSfGFP::LPR-3]) X | This work |
| UP4382 | bli-6(ju1914 [BLI-6::mNG::3xFLAG]) IV; lpr-3(cs266[mCherry::LPR-3]) X | This work |
| UP4435 | lpr-3(cs266 cs315 [mCherry::LPR-3(ΔNter)]) X | This work |
| UP4437 | wrt-10(syb7022[WRT-10::sfGFP])II; lpr-3(cs266 cs315 [mCherry::LPR-3(ΔNter)])X | This work |
| UP4441 | lpr-3(cs266 cs315 [mCh::LPR-3(DNter)]) X; csEx436 rescue | This work |
| UP4464 | bli-6(ju1914 [BLI-6::mNG::3xFLAG]) IV; lpr-3(cs266 cs315 [mCherry::LPR-3(ΔNter)]) X; csEx436 [lpr-3 fosmid + myo-2::mCherry] | This work |
| UP4465 | wrt-10(aus36); bli-6(ju1914 [BLI-6::mNG::3xFLAG]) IV | This work |
| UP4476 | wrt-10(aus36) II; lpr-3(cs266 cs315 [mCherry::LPR-3(ΔNter)]) X | This work |
| UP4477 | wrt-10(aus36) II; bli-6(ju1914 [BLI-6::mNG::3xFLAG]) IV; lpr-3(cs266 cs315 [mCherry::LPR-3(ΔNter)]) X | This work |

**S2 Table.**
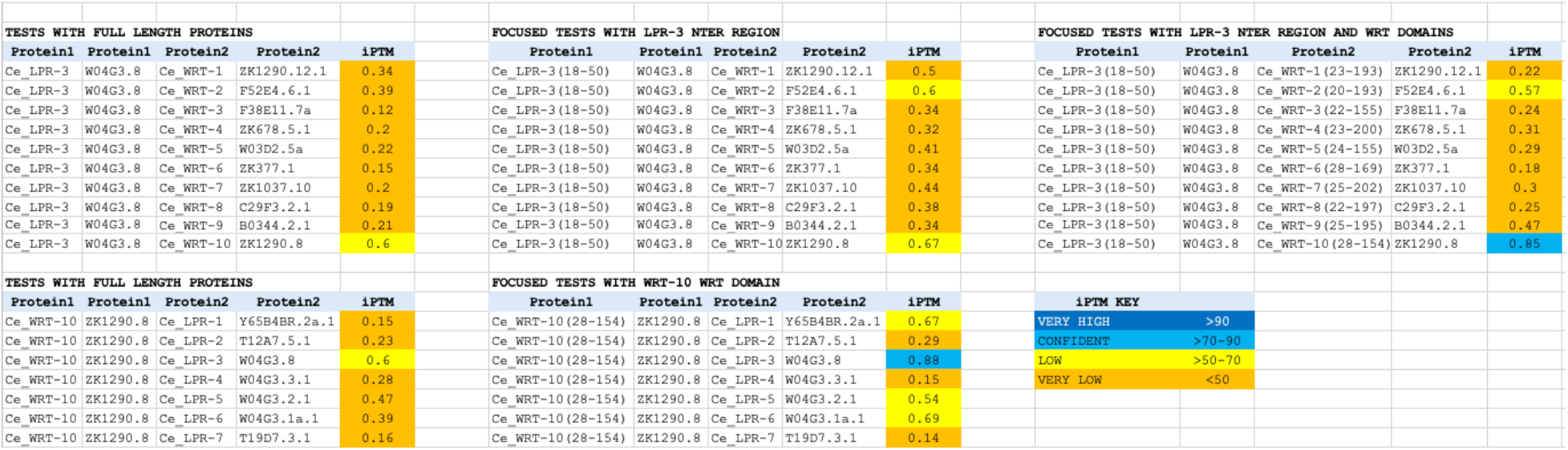
Alphafold3 pairwise test results of *C. elegans* lipocalins and WRT domain proteins.

**S3 Table.** Source data for Fig. 3C.

**S4 Table.** Source data for Fig. 4B.

**S5 Table.** Source data for Fig. 4D.

**S6 Table.** Source data for Fig. 5C.

**S7 Table.** Source data for Fig. 5E.

**S8 Table.** Source data for S2 Fig. panel C.

**S9 Table.** Source data for S2 Fig. panel F.

**S1 Fig.**
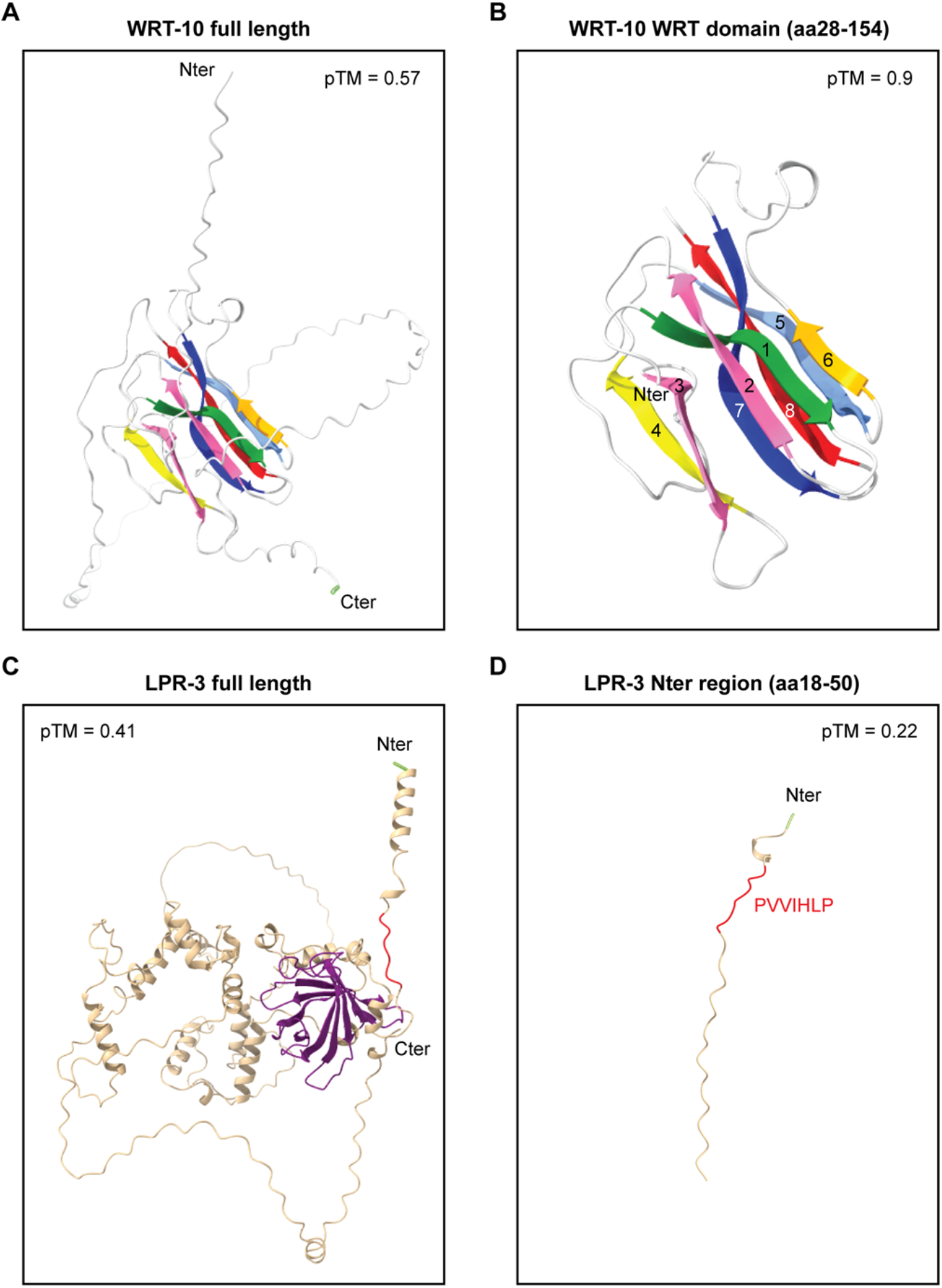
Alphafold3 predicted structures for WRT-10 and LPR-3. **A.** Full-length WRT-10. **B.** WRT-10 WRT domain (amino acids 28-154). Predicted β-strands are colored and numbered beginning at the most N-terminal region. **C.** Full length LPR-3. **D.** LPR-3 N-terminal region (amino acids 18-50). The region predicted to bind WRT-10 (PVVIHLP) is indicated in red. Alphafold3 pTM scores are indicated.

**S2 Fig.**
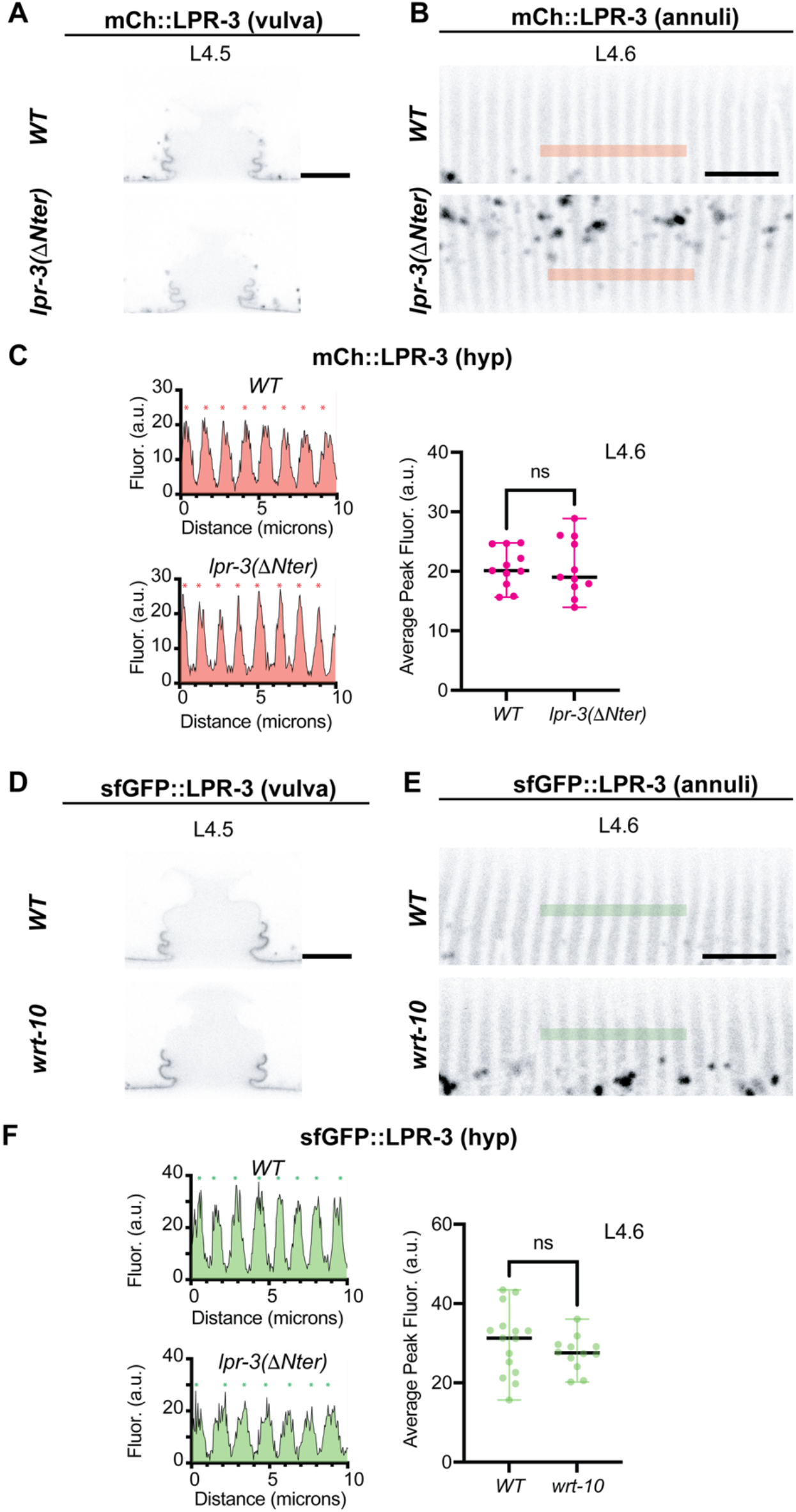
*lpr-3(6.Nter)* and *wrt-10(-)* mutants do not disrupt LPR-3 patterning in the developing vulva or hyp7 annuli. **A,B,D,E.** mCh::LPR-3 (A, B) or sfGFP::LPR-3 (D, E) in developing vulva (A,D) and annuli (B,E) in the indicated genotypes and stages, shown in inverted grayscale. Images in B and E are maximum intensity projections of three confocal z-slices. Colored bars in B,E indicate method for quantifying fluorescence intensity in C,F. Scale bars, 5 microns. **C,F.** Quantifications of peak fluorescent signal intensity across the L4.6 annuli as measured using the FIJI Plot Profile tool (n>10 for each genotype). Example line scans are shown at left and asterisks indicate called peak regions. See Materials and Methods for additional details. ns, not significant, Mann-Whitney U test.

**S3 Fig.**
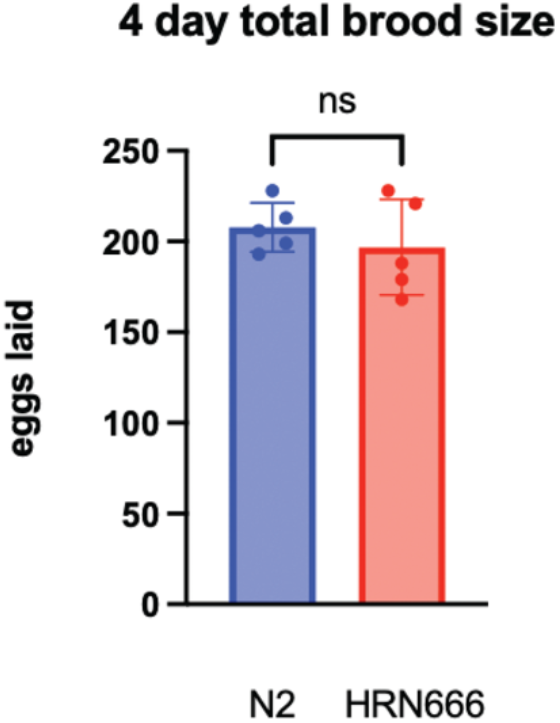
*wrt-10(-)* mutants have normal brood size. Fertilized eggs released by five N2 (WT) and five HRN666 (*wrt-10(aus36))* self-fertilizing hermaphrodites were monitored from lay initiation to cessation on day 4. ns, no significant difference, Mann-Whitney U test. Note that this experiment was not powered to detect subtle changes, nor did we assess broods from mated animals.

## Notes

### Competing Interest Statement

The authors have declared no competing interest.

